# AUTS2 gene dosage affects synaptic AMPA receptors via a local dendritic spine AUTS2-TTC3-AKT-mTORC1 signaling dysfunction

**DOI:** 10.1101/2022.12.01.518705

**Authors:** Aude-Marie Lepagnol-Bestel, Arnaud Duchon, Julia Viard, Mirna Kvajo, Rachel Daudin, Malik Khelfaoui, Simon Haziza, Yann Loe-Mie, Mattia Aime, Futoshi Suizu, Marie-Christine Birling, Mounia Bensaid, Sylvie Jacquot, Pascale Koebel, Céline Reverdy, Jean-Christophe Rain, Masayuki Noguchi, Xavier Marquez, Antoine Triller, Yann Humeau, Yann Hérault, Maria Karayiorgou, Joseph A. Gogos, Michel Simonneau

## Abstract

The Human 1.2-Mb *AUTS2* locus on chromosome 7q11.22 encodes a 1259-aa full-length protein, and a 711-aa C-terminal isoform. Functions of these AUTS2 proteins are only partly known. The major traits found in patients displaying *AUTS2* locus mutations are Intellectual Disabilities, microcephaly attention deficit hyperactivity disorder (ADHD) (54%), and autistic traits. Furthermore, *AUTS2* common variants were recently found associated to alcohol consumption and dyslexia using GWAS approaches. Auts2 localizes mainly in cell nuclei. We evidenced by super-resolution that Auts2 is present in dendritic spines. Auts2 interacts with Ttc3, the Akt2 E3 ligase, and negatively regulates Akt2 ubiquitination. Auts2 haploinsufficiency affects Akt/mTorc1 pathway with a decrease in AMPA and NMDA receptor subunits and in synaptic currents. Akt2 injection in postsynaptic neurons is sufficient to reverse changes in synaptic currents generated by Auts2 haploinsufficiency. Using chromosome engineering based on targeted meiotic recombination, we generated two mouse models with *Auts2* locus deletion and duplication. Deleted *Auts2* locus mice display stereotypies (rearing), perseveration and abnormal recognition memory. Duplicated *Auts2* locus mice display similar perseveration and abnormal recognition memory but also a decrease in cued and contextual fear memory. Gene dosage induce changes in brain sub-region neuronal networks. In the thalamo-lateral amygdala pathway linked to cued fear memory, we found synaptic impairments linked to AMPA receptors, with a specific decrease in pAKT/total AKT ratio in duplicated Auts2 mice. Altogether, our study thereby provides a novel mechanistic and potentially therapeutic understanding of synaptic AKT/mTORC1 deregulated signaling and its related behavioral and cognitive phenotypes.

---

Autism spectrum disorder (ASD) is a broad group of brain development disorders characterized by impaired social interaction and communication, repetitive behavior and restricted interests (1). The genetic architecture of ASD involves the interplay of common and rare variants and their impact on hundreds of genes (2). One of promising ASD candidate genes is Autism susceptibility 2 (AUTS2) (3). Number of studies have reported multiple genetic variations in (*AUTS2*) that are associated with autism and related neurodevelopmental disorders (NDDs) since the identification of *Autism susceptibility candidate 2* (*AUTS2*) gene translocation in a pair of autistic twins (4). These variants have been found in ASD, mental retardation, epilepsy and a variety of psychiatric diseases (3, 5, 6). *AUTS2* common variants were recently found associated to alcohol consumption and dyslexia using GWAS approaches (7–9). Furthermore, both complete duplication and deletion of ~1Mb *AUTS2* locus has been reported as giving similar ASD phenotypes (10). However, the influences of *AUTS2* dysfunction on ASD-related behaviors, as well as underlying developmental and neural circuit mechanisms, are still unclear. A possible involvement of AUTS2 in synaptic function as for many risk genes of ASDs (2, 11, 12) was not yet investigated. Our working hypothesis was that AUTS2 takes part in synaptic protein complexes with functional impact that varies with the dosage of protein partners, in order to explain the variety of behaviour and cognitive subtypes found in AUST2 patients. Here, we revealed a previously unidentified localization of AUTS2 in dendritic spines and the interaction of AUTS2 on TTC3 that is involved in the AKT/mTORC1 pathway in dendritic spines, leading to a decrease of synaptic AMPA receptors. We generated novel deletion and duplication Auts2 models of deletion and duplication of the ~1Mb *AUTS2* locus using genomic engineering in order to identify ASD-related behaviors in these mice, to analyze ASD-related behaviors and to analyze possible synaptic defects in involved neural circuits.

## Auts2 is expressed in dendritic spines and negatively regulates Akt2 degradation via its interaction with Ttc3

In spite of many mouse Auts2 models that were generated, only partial subcellular and molecular information is available on mechanisms of AUTS2 in normal and pathological brain development (13, 14).

Previous studies showed that AUTS2 localizes both in nuclei and in cytoplasm (15). In nuclei, AUTS2 associates with a polycomb repressive complex 1 (PRC1) and activates genes in the developing brain (16, 17). AUTS2 is also part of a Rac1 complex to control actin dynamics in neuronal migration and neuritogenesis (15). However, no studies evidenced yet a localization of AUTS2 protein at the synapse.

We first studied a possible synaptic localisation of Auts2 protein using analysis of mouse brain subcellular fractions. We detected Auts2 protein in both nuclear fractions and synaptosomes isolated from mouse adult brain (Fig. 1A). Using confocal microscopy of mouse primary hippocampal cultures at day in culture (DIC) 21, we found that Auts2 was localized in discrete puncta overlapping with two dendritic spine proteins, post synaptic density 95 (PSD95) localized in dendritic spine head and synaptopodin (SYNPO) localized in dendritic spine neck (Fig. 1B; Fig. S1).

**Fig.1.**
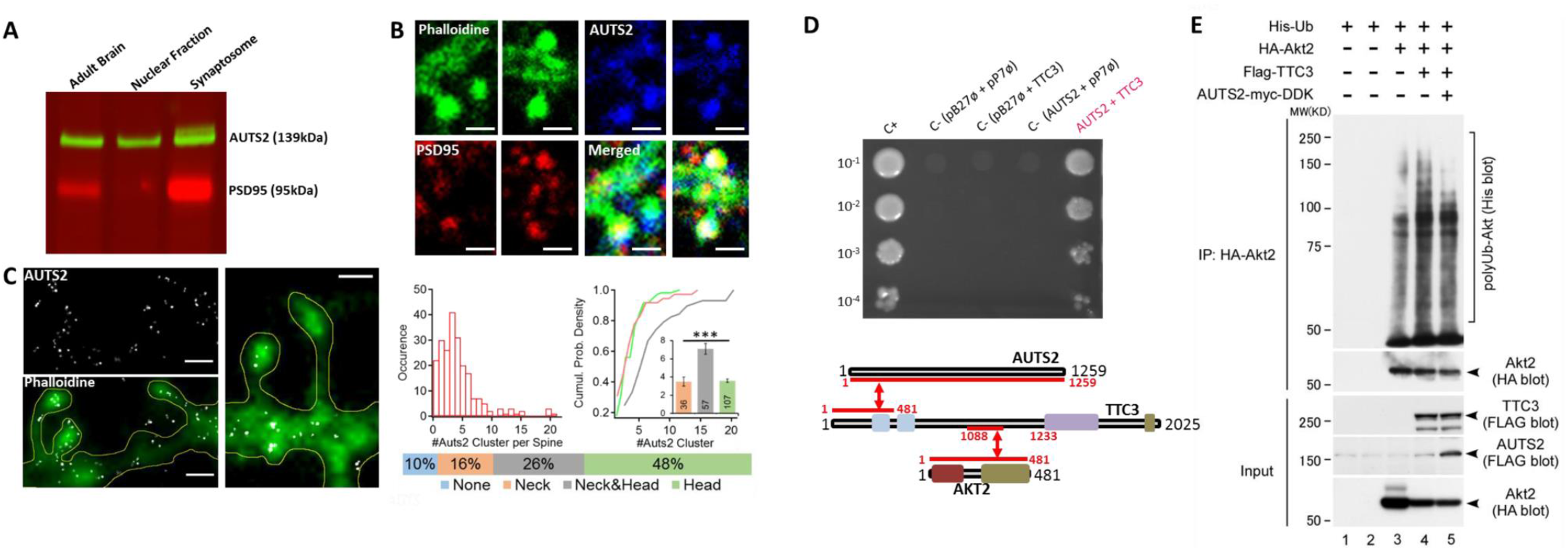
Super-resolution localization of Auts2 in dendritic spines and AUTS2 as an inhibitor of AKT2 E3 ligase,TTC3. **A.** Fluorescent immunoblot expression of Auts2 protein (green band) in adult mouse brain, adult mouse brain nuclear enrichment fraction and adult mouse cortical synaptosomal enrichment fraction. Enrichment purity was controlled with PSD95 protein expression (red band). **B.** Confocal imaging of immunocytochemistry on DIC21 primary hippocampal neurons using AUTS2 antibody and the postsynaptic marker PSD95. Enlargements of dendritic spines images are shown. Scale bar=10μm. **C.** Super-resolution dSTROM imaging of immunocytochemistry on DIC21 primary cortical neurons using AUTS antibody (white dots) and conventional imaging of phalloïdine (green). Histogram representing Auts2 cluster per dendritic spines, with in average 4,1±0.3 clusters/spine. Cumulative probability density of the number of Auts2 cluster in neck (salmon), neck&head (purple) and head (green). Inset: bar plots show the average of Auts2 cluster per spine sub-region, with 3.6±0.2 clusters in the head, 3.5±0.5 clusters in the neck and 7.1±0.6 clusters in the entire spine (neck&head). Figures inside bar plots represent the number of samples. Localization of Auts2 clusters inside the spine head (green), the neck (salmon) or both of them (grey). Statistical significance was calculated using the Mann-Whitney test (***: p<0.001). Scale bar=1μm. **D**. Yeast two-hybrid one-by-one assays revealed TTC3 as an interactor of AUTS2. Schematic representation (lower panel) of the AUTS2, TTC3 and AKT interactions demonstrated in the study. C+: interaction positive control; C- (pB27ø + pP7ø): empty pB27 vector + empty pP7 vector; C- (AUST2 + pP7ø): pB27-AUST2 + empty pP7 vector; C- (pB27ø + TTC3): empty pB27 vector + pP6-TTC3; AUST2 + TTC3: pB27-AUST2 + pP6-TTC3 **E**. AKT2 polyubiquitination by TTC3 is reduced by AUTS2 expression in HeLa cells. TTC3 enhanced AKT ubiquitination (lanes 3 vs 4 in top panel), however, ectopic AUTS2 expression reduced Akt ubiquitination by TTC3 (lanes 4 vs 5 in top panel).

We analysed sub-compartmentalization of Auts2 clusters both inside dendritic shafts and spines using dSTORM high resolution microscopy (Fig. 1C). Half of the dendritic spines have clusters only inside spine heads and only 10% of them show no Auts2 clusters. Surprisingly, Auts2 clusters are preferentially localized in the neck for SYNPO-positive spines and in the head for SYNPO-negative spines (Fig. S2).

We next searched for protein interactors using yeast two-hybrid (Y2H) screens (18) and identified that AUTS2 binds tetratricopeptide repeat domain 3 protein (TTC3) which acts as ubiquitin E3 ligase specific for AKT (19). We confirmed the interaction between TTC3 and detected two distinct TTC3 binding domains in AUTS2 using the one-by-one Y2H assay (Fig. S3A-B). AUTS2 interacts with TTC3 in a domain located to the N-terminal part of the protein (1-481 amino acid domain) that includes the first tetratricopeptide repeat (TPR) motif and S378, a key AKT-dependent phosphorylation site (19) (Fig. 1D). Furthermore, we evidenced Auts2-Ttc3 and Ttc3-Akt2 direct interactions in dendrites and dendritic spines of hippocampal neurons (Fig. S4), using the *in situ* proximity ligation assay (PLA), which enables detection of endogenous protein-protein interaction events (20). We found that the three Akt (Akt1, Akt2 and Akt3) are present in synaptosomes (Fig. S4-A). However, we detected a similar number of interactions for Ttc3-Akt2 as compared to Ttc3-panAkt, suggesting that Akt2 interacts only with Ttc3, in dendritic spines of primary cultures of mouse hippocampal neurons (Fig. S4-B).

It was already reported that TTC3 facilitated AKT degradation (19), suggesting that the size of the AKT pool in a subcellular compartment can be modulated via TTC3-mediated degradation. Therefore, AUTS2 downregulation may induce an increase in TTC3 E3 ubiquitin ligase activity, leading in turn to a decrease of the AKT pool. To test this hypothesis, we studied AKT2 polyubiquitination induced by TTC3 in presence or absence of AUTS2, in HeLa cells. AUTS2 reduced TT3-induced AKT2 polyubiquitination in HeLa cells (Fig. 3E). This effect was confirmed in 293T cells with two AKT2 mutants: AKT2 mutant 1 (AKT2-Glu17Lys) and constitutively active AKT2 mutant 2 (AKT2-T309E-S474D) (Fig. S3D).Taken together these results demonstrate that AUTS2 negatively regulates AKT degradation via its interaction with TTC3.

**Fig. 2.**
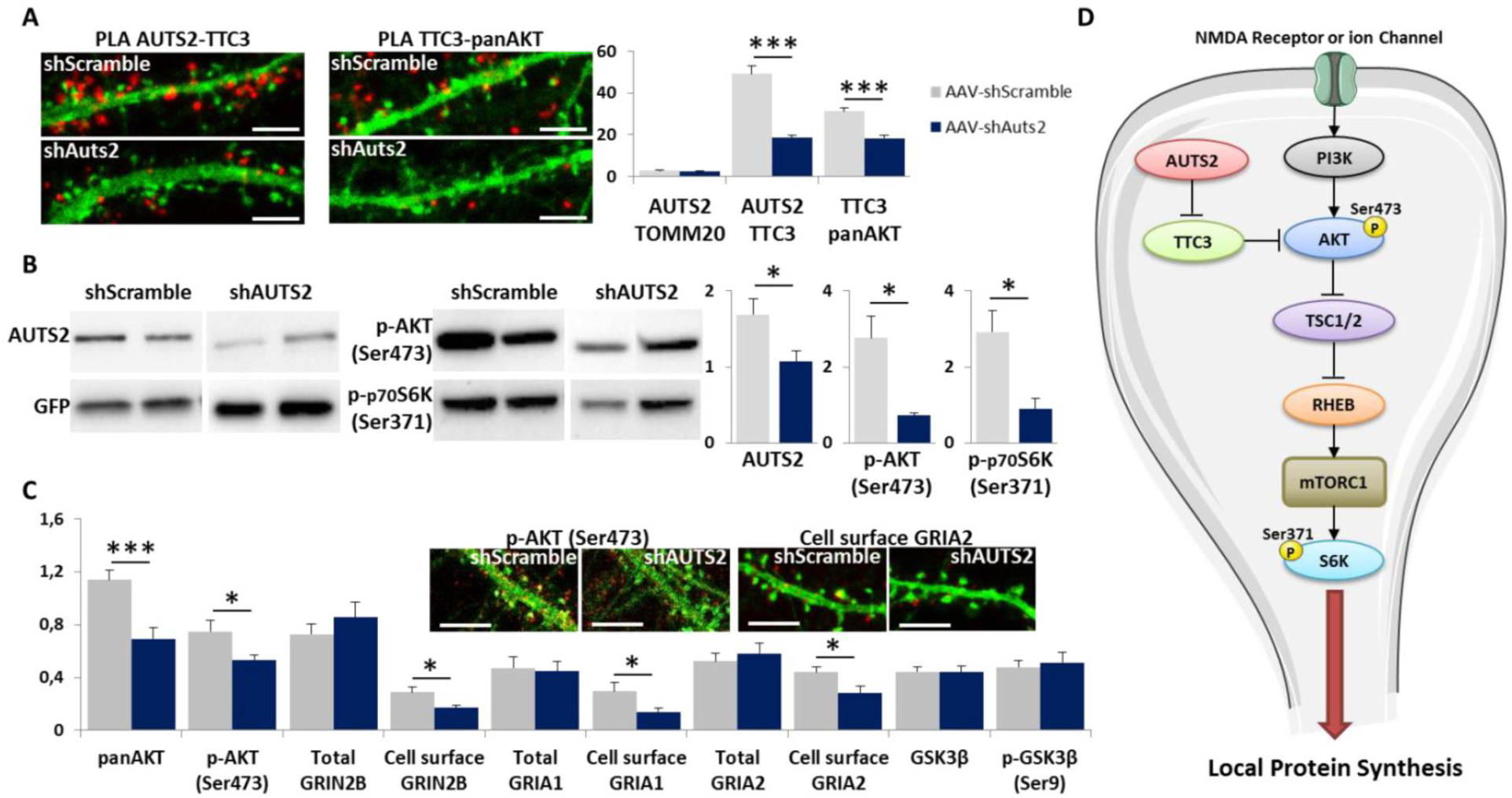
Characterization of an Auts2/Ttc3/Akt/mTORc1/S6K pathway impaired in AAV Auts2-silenced cortical primary neurons. A. In situ proximity ligation assays PLA (left panel) on primary cortical neurons fixed at DIC21 (red fluorescence) using anti-Auts2 and anti-Ttc3 antibodies, anti-Ttc3 and anti-panAkt antibodies as a positive control and anti-Auts2 and anti-Tomm20 antibodies as a negative control. Immunohistochemistry was performed using anti-Phalloidine (green fluorescence) to visualize dendrites and dendritic spines. Scale bars=10μm. Mean PLA interaction point numbers (right panel) were calculated in 30 dendrites of 150μm under control (AAV-Scramble) and Auts2 knockdown (AAV-shAUTS2) conditions. *** p < 0.0005 B. Immunoblotting of Auts2, phospho-Akt i.e. p-AKT (Ser473) and phospho-p70S6K i.e. p-p70S6K(Ser371) on primary cortical neurons fixed at DIC21 under control (AAV-Scramble) and Auts2 knockdown (AAV-shAUTS2) conditions. Quantification of each protein was performed on at least 3 independent samples for each condition and expression was normalized against the GFP protein. *p<0.05 C. Immunocytochemistry of phospho-Akt i.e. p-AKT (Ser473) and cell surface GRIA2 receptors (red fluorescence) on primary cortical neurons fixed at DIC21 under control (AAV-Scramble) and Auts2 knockdown (AAV-shAUTS2) conditions. Scale bars=10μm. Mean point numbers were calculated in at least 13 dendrites of 100μm for each condition and expression was normalized against the green area. ** p < 0.005. D. Schematic representation of AUTS2-TTC3-AKT molecular pathway in dendritic spine.

**Fig. 3.**
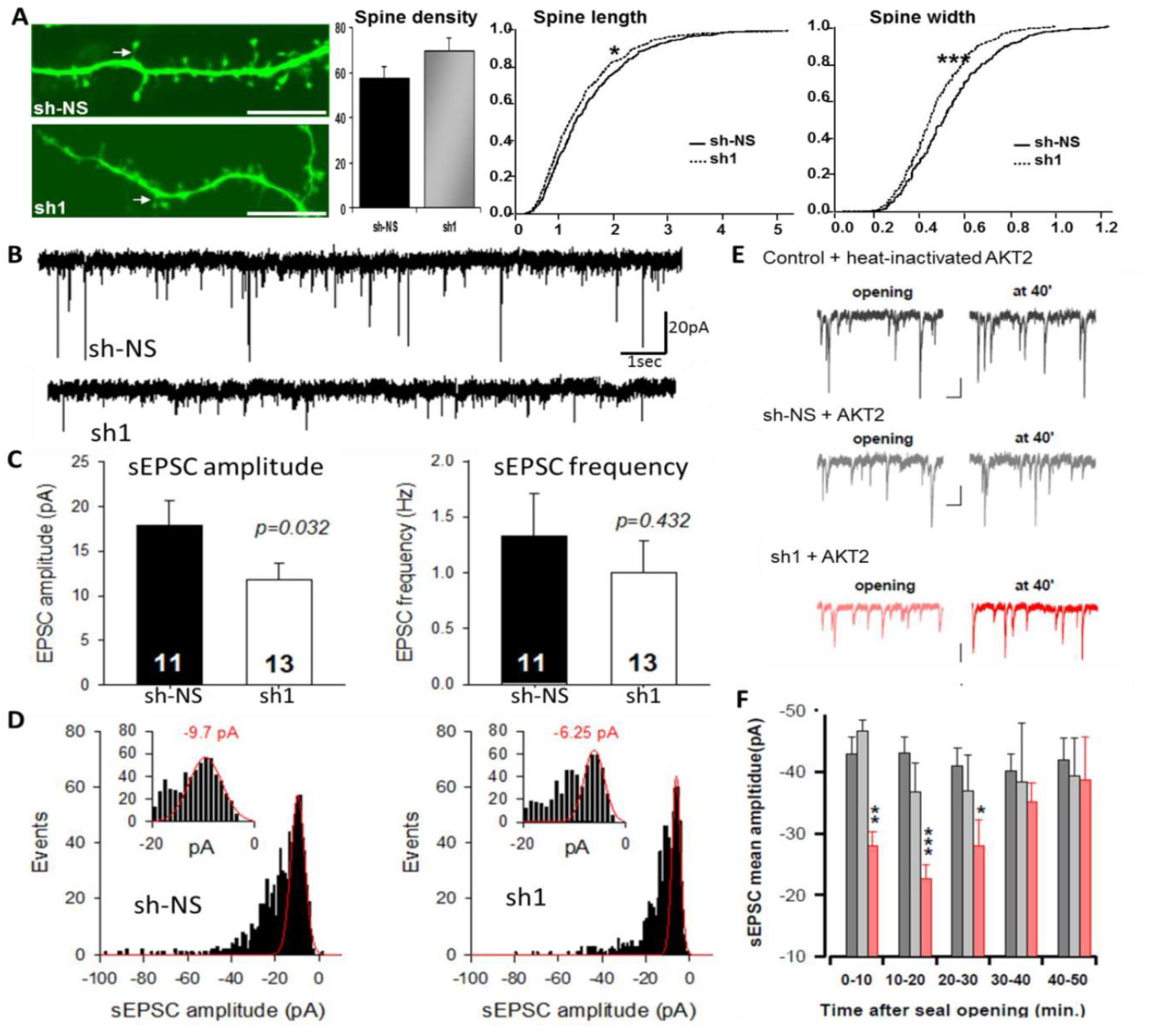
Impaired synaptic functions in Auts2-silenced cortical primary neurons and rescue of synaptic currents by AKT2 postsynaptic intracellular injection. A. Dendritic spines from E17.5 primary cortical neurons transfected at DIC7 with sh-NS or sh1 and fixed at DIC21. Mushroom spines are indicated with arrows. Scale bars=10μm. Total dendritic spine number was calculated per 75 μm dendritic fragments. Dendritic spine length and spine width are calculated respectively as the length of the neck and the width of the synaptic button of the mushroom spines present on 75 μm dendritic fragment samples. *p<0.01; **p<0.001; ***p<0.0001 B. Examples of whole-cell voltage-clamp recordings of sEPSC recorded in post-synaptic cortical neurons transfected either with sh-control (sh-NS) or sh-Auts2 (sh1). C. Quantification of whole-cell voltage-clamp recordings sEPCS amplitudes and frequencies. Note that silencing of Auts2 induced a significant decrease in sEPSC amplitude without significant change in sEPSC frequency. D. Quantal distribution of whole-cell voltage-clamp recordings sEPCS of control as compared to Auts2-silenced post-synaptic cortical neurons. An enlargement of the distribution of sEPSC for the lowest quantal release is inserted for both control and Auts2-silenced postsynaptic cortical neurons. Note that this quantal component amplitude decrease from −9.7 to −6.25 pA suggesting a post-synaptic effect of Auts2 silencing. E: Sample whole-cell voltage-clamp recordings of sEPSC recorded in post-synaptic cortical neurons following heat-denatured or native Akt2 injection. F. Quantification of sEPCS amplitudes as a function of time following seal openings. Note the recovery of sEPCS 40 minutes after Akt2 injection.

## Down-regulation of Auts2 expression induces cellular and molecular dendritic spine changes linked to AKT-mTORC1 pathway

The AUTS2 regulation of AKT is of particular interest as AKT kinases regulate dendritic spine morphogenesis (21, 22). Furthermore, AKT/mTORC1 pathway was already involved in the pathophysiology of ASD for TSC1/TSC2 (23) and for Shank3 (24).

We investigated possible functional changes in the AKT/mTORC1 pathways (25) using of AAV Auts2 shRNA (shAuts2) to mediate gene knockdown in primary mouse neurons. We first quantified the dendritic spine density and morphology *in vivo* using shAuts2 stereotaxic injection in CA3 hippocampal neurons of adult mice. We quantified a decrease in mushroom spine density with a significant increase of their length and width (Fig. S5).

Using primary cultures of cortical neurons, we evidenced a decrease in the number of Auts2-Ttc3 and Ttc3-Akt complexes upon Auts2 depletion in dendritic shafts and spines (Fig. 2A). We also found a significant decrease in the quantity of Akt and phospho-Akt as well as a significant decrease of the ratio of phospho-Akt/Akt in *Auts2*-silenced neurons, using western blotting (Fig. 2B).

We next analysed the phosphorylation of key proteins in the AKT/mTORC pathway. mTOR protein is a large protein kinase that nucleates at least in two distinct multi-protein complexes i.e. mTORC1 and mTORC2 which involves separate molecular pathways (25). Phospho-p70s6k is a downstream effector of mTORC1 leading to protein translation and Gsk3β is a downstream effector of mTORC2 leading to metabolism. We found a decrease of phospho-p70s6k and no change in p-GSKβ upon Auts2 depletion, indicating a specific impact on mTORC1 pathway (Fig. 2B). We hypothesized that *Auts2*-silencing led to a decrease in local protein translation, in particular membrane glutamate receptors Gria and Grin2b known to be key proteins in synaptic function. Using immunocytochemistry, we confirmed the local decrease of Akt and phospho-Akt in dendritic shafts and spines in *Auts2*-silenced neurons. Furthermore, we showed a local decrease in cell membrane Gria1, Gria2 and Grin2a receptors at the dendritic shafts and spines under Auts2 knockdown whereas neither Gsk3β, phospho-Gsk3β nor any of the total receptors did locally change (Fig. 2C). Taken together, these results demonstrate that Auts2 induces a decrease of the Akt and phospho-Akt pool at the dendritic spine leading to a decrease of both Gria1 and Grin2b glutamate receptors localized at the neuronal membrane of dendrites and dendritic spines (Fig. 2D).

## Down-regulation of Auts2 expression induces changes in synaptic function

We next investigated if *Auts2* knockdown affects synaptic function using primary neurons transfected with Auts2 shRNA vectors (Fig. 3). We evidenced a significant increase in the total number of dendritic spines and a significant decrease of the mushroom spine length and width under *Auts2* knockdown (Fig. 3A). Such morphological changes in mushroom spines are linked to abnormal spine maturation and synaptic function. These changes found in primary neurons are distinct from those we report in slices of *Auts2* knockdown hippocampus (Fig. S5), suggesting distinct modifications in the Rac1 pathway (15). We next analysed synaptic impairment and found that spontaneous excitatory postsynaptic current (sEPSC) amplitude responses were impaired in *Auts2* silenced postsynaptic neurons (Fig. 3B-C). In contrast, frequencies of sEPSC were not modified (Fig. 3C). Furthermore, quantal analyses of sEPSC responses indicate a decrease in the elementary event, suggesting that *Auts2* silencing in postsynaptic neurons induce a decrease in the elementary postsynaptic response that can be attributed to changes in properties of postsynaptic receptors (Fig. 3D). Taken together these data strongly suggests that *Auts2* deficiency may affect ultimately synaptic connectivity leading to abnormal function of excitatory synapses.

## AKT2 postsynaptic intracellular injection rescue impaired synaptic functions

We hypothesized that the observed decrease in synaptic current amplitude induced by Auts2 depletion was due to a local depletion of AKT molecules. As we found that only Akt2 interacts with TTC3 in dendritic spines (Fig. S4), we propose that depletion of Akt2 in dendritic spines is sufficient to induce impaired synaptic functions. We tested this hypothesis with AKT2 injection into *Auts2*-deficient cortical neurons and succeed in rescuing the altered synaptic phenotype. Introduction of nanomolar concentrations of AKT2 into *Auts2*-deficient neurons reversed the size of miniature postsynaptic currents to wild-type levels in a time period compatible with the trafficking of AKT2 from the soma to dendritic spines (Fig. 3E-F). Taken together, these results indicate that functional synaptic defect can be rescued by AKT2 postsynaptic injection.

## Generation of mice with a complete deletion or duplication of the Auts2 locus

All mouse models available so far do not imply a deletion of all isoforms and none of them is based on a mutation of *AUTS2* gene detected in patients (13, 14). As both complete duplication and deletion of ~1Mb *AUTS2* locus has been reported as giving similar ASD phenotypes (10), we generated mouse lines with a complete deletion or duplication of the *Auts2* locus. The human *AUTS2* and the mouse *Auts2* loci are highly homologous and span ~1Mb.

Deletion and duplication of *Auts2* locus uses the trans allelic targeted meiotic recombination (TAMERE) strategy (26). This strategy is well adapted for large transcription units, clustered genes and chromosomal loci that require the design of novel experimental tools to engineer genomic macro-rearrangements. This protocol allows for the combination of various alleles within a particular locus as well as for generation of interchromosomal unequal exchanges (Fig. S6). LoxP in the 5’ and the 3’ of the *Auts2* locus were introduced in ES cells. The exact position of the deletion and the duplication on mouse chromosome 5 is between 131,427,079 and 132,557,948 (GRCm38/mm10).The size of the deleted or duplicated fragment is 1.13 Mb. The deletion Del(*Auts2*)1Ics and the corresponding duplication Dp(*Auts2*)2Ics, that will be noted here Del/+ and Dup/+, were established on a pure C57BL/6N genetic background. Auts2 expression levels in hippocampus of adult Del/+ and Dup/+ mice reveal a decreased and an increase in expression respectively, as compared to control littermates (Fig. 4A; Fig. 5A).

**Figure 4:**
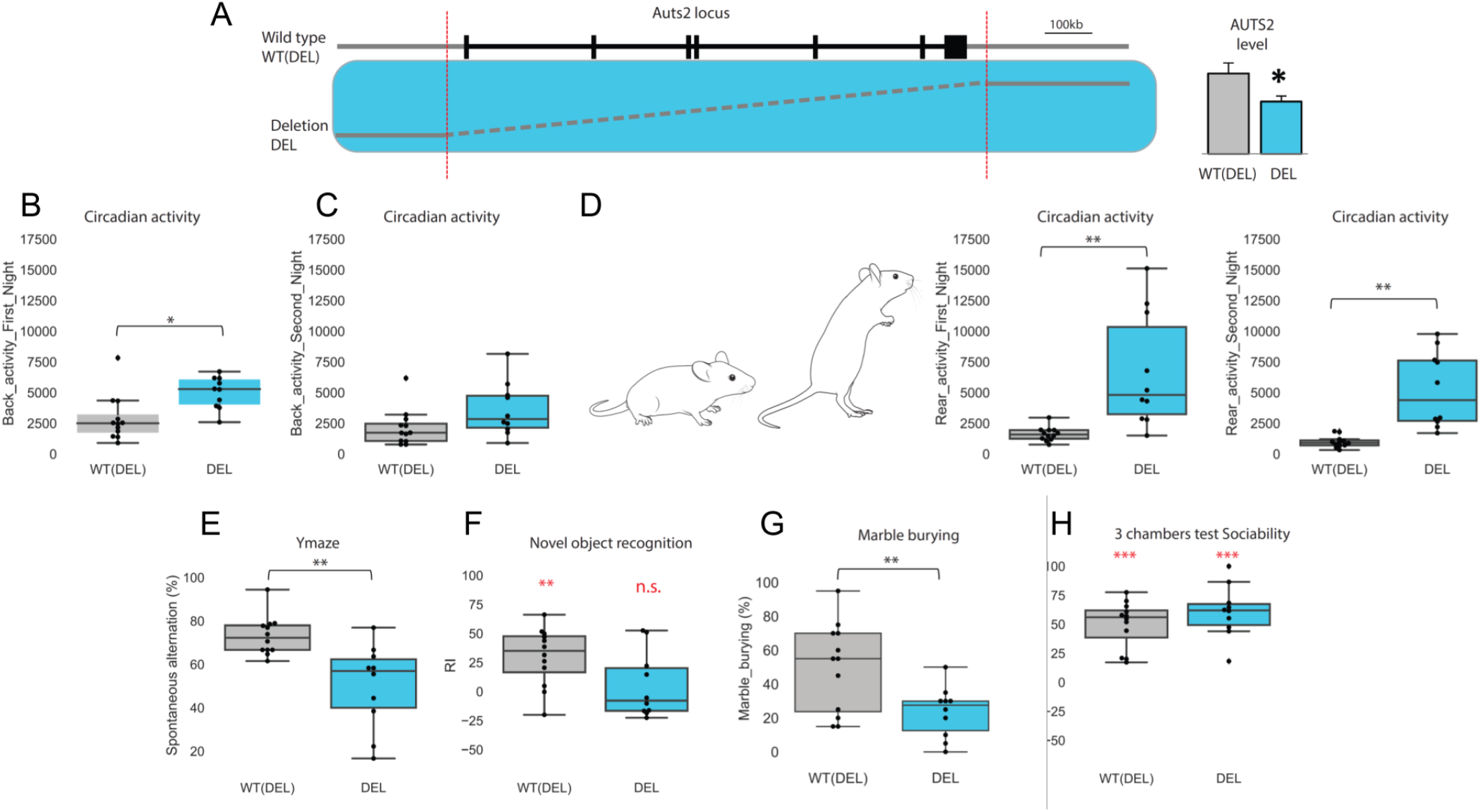
Del/+ Auts2 behavioural impairments. **(A)** Schematic representation of the deletion produced by TAMERE strategy. Validation of Auts2 protein expression in control WT (DEL) compared to transgenic (DEL) adult hippocampus was performed from at least 4 transgenic mice. Protein expression was normalized using β-Actin protein. Auts2 expression level reveal a decreased expression in DEL mice. (values represent means+/− SEM). **(B-D)** During the circadian activity test, the DEL mice were hyper-active with an increased back activity **(B,C)** or rear activity (or stereotypies) during the first and second night **(D)**. **(E)** The spontaneous alternation rate in Y-maze during a single 8-min session indicates a lower percentage of alternation for the DEL mice compare to the control littermate animals. **(F)** In the novel object recognition test, DEL mice show a recognition memory deficit after a retention of 24 hours with no discrimination between the familiar and the novel object compared to the WT (DEL). The one sample Ttest indicated that RI was not significantly different from 0, corresponding to the chance level. **(G)** In the marble burying test, the percentage of marble buried is decrease in the DEL compare to control littermate**. (H)** The social interaction test, with the recognition index, does not unravel change in DEL mice compare to their littermates either during the sociability phase or during the social discrimination phase (not show). Box plots with the median and quartiles. * p<0.05. **p<0.01. ***p<0.001. One sample T test result versus 0 was indicated in red colour in the graphic for RI.

**Fig. 5.**
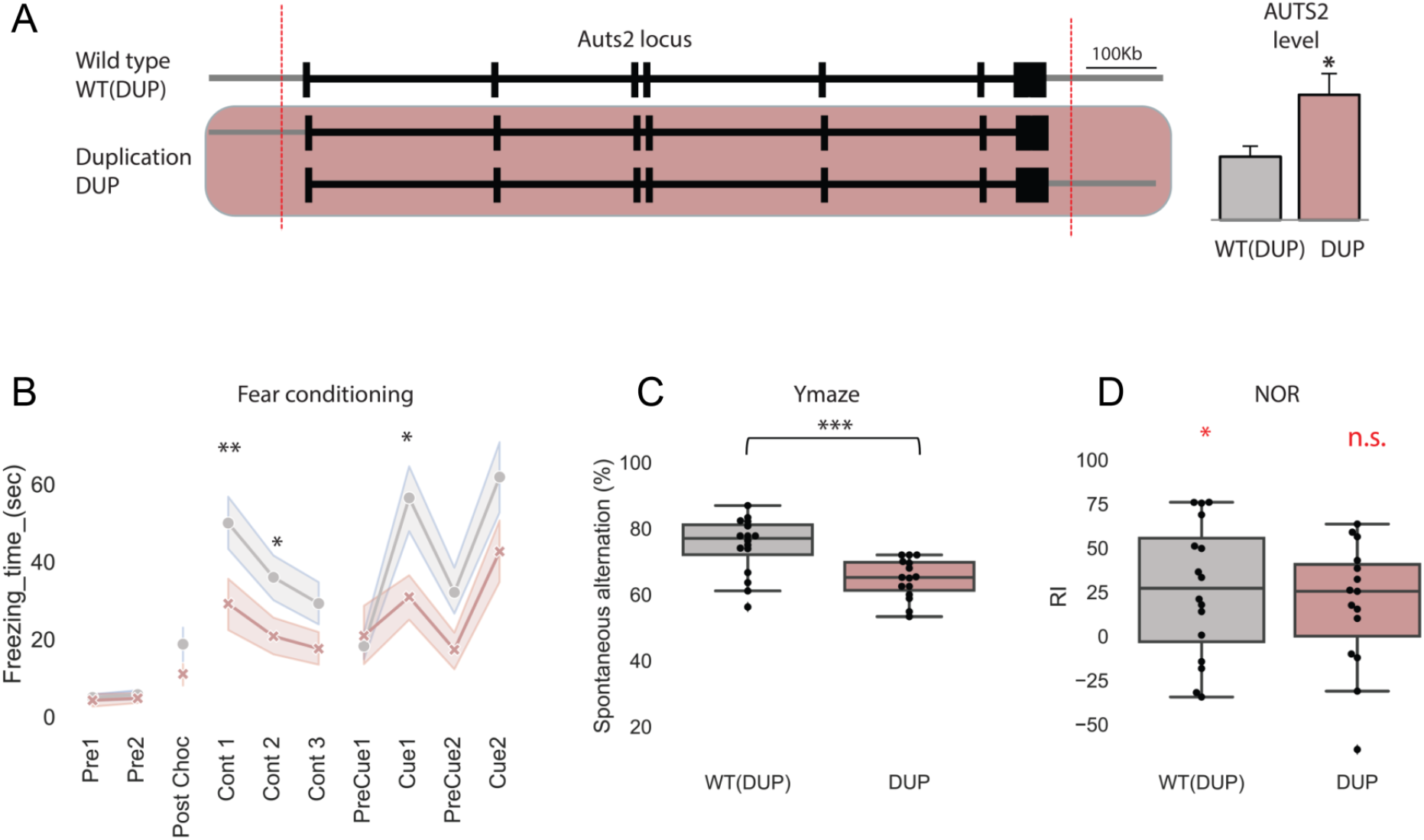
Dup/+ Auts2 behavioural impairments. **(A)** Schematic representation of the duplication produced by TAMERE strategy. Validation of Auts2 protein expression in control WT (DUP) compared to transgenic (DUP) adult hippocampus was performed from at least 4 transgenic mice. Protein expression was normalized using β-Actin protein. Auts2 expression level reveal a increased expression in DUP mice. (values represent means+/− SEM. *P < 0.05). DUP mice are impaired in the contextual and cued fear conditioning, illustrated by a decreased freezing time (sec) in both test session **(B)**. Alternation rate in Y-maze during a single 8-min session indicates a lower percentage of alternation for the DUP mice compare to the WT (DUP) **(C)**. DUP animals show a recognition memory deficit after a retention of 24 hours with no significant (ns) discrimination between the familiar and the novel object (RI not significantly different from 0) **(D)**. Box plots with the median and quartiles. * p<0.05. **p<0.01. ***p<0.001. One sample T test result versus 0 was indicated in red colour in the graphic for RI.

## Deletion and Duplication of Auts2 lead to autism-related behaviors in mice

To study the potential effect of *Auts2* locus deletion on ASD-related behaviors, we conducted a battery of tests (Fig. 4B-H; Fig.S7; Fig S8). We explored first repetitive and restricted behaviors are one of the core features of ASD (27). We evidenced hyperactivity in Del/+ mice and rearing behavior (Fig. 4C-D). We found perseveration in the Y-maze for Del/+ mice. Furthermore, object interactions of Del/+ mice is impacted for both Novel Object Recognition (NOR) test (Fig. 4E) and marble-burying test (Fig. 4G). NOR impairment is an indicator changes in perirhinal-lateral entorhinal cortex pathway (28). Decrease in marble-burying test can be also considered as an avoidance behavior indicative of a decrease in anxiety (29). In contrast, we found no changes in social interactions of Del/+ mice as analyzed by the three-compartment chamber test (Fig. 4H).

Similar behavioral tests were used for Dup/+ mice (Fig. 5B-D; Fig.S7; Fig S8). Dup/+ mice showed abnormalities during either contextual or cued fear compared to WT littermates (Fig. 5B). ASD display prominent social behavior abnormalities that are known to involve diverse brain microcircuits including amygdala (30–32). Furthemore, we evidenced changes in Y-maze and NOR similar to that described in Del/+ mice (Fig. 5C-D).

## Impaired synaptic function in thalamo-lateral amygdala of Dup/+ Auts2 mice

We took advantage of Dup/+ Auts2 mice to elucidate molecular, cellular, and neural circuit changes that occur in the brain during learning fear conditioning (33). Dup/+ Auts2 mice display abnormalities during either contextual or cued fear conditioning compared to their WT littermates (Fig. 5B). In this form of learning, animals associate two stimuli, such as a tone and a shock. This auditory fear conditioning is a well-characterized behavioural paradigm in which an animal learns to associate a tone with an electric shock and subsequently freezes when presented with a tone alone (34). Molecular mechanisms have been elucidated (35). Fear conditioning drives AMPA type glutamate receptors into the synapse of a large fraction of postsynaptic neurons in the lateral amygdala (LA), a brain structure essential for this associative learning process. Furthermore, memory was reduced if AMPA receptor synaptic incorporation was blocked in as few as 10 to 20% of LA neurons. Thus, the encoding of memories in the lateral amygdala is mediated by AMPA receptor trafficking.

Our working hypothesis was that the thalamo-LA excitatory synapses that use AMPA receptors were affected by *Auts2* duplication. To test this hypothesis, we recorded thalamo-lateral amygdala synapses in coronal acute slices of adult wild type (WT) and Dup/+ *Auts2* (DUP) mice (Fig. 6A). The amplitude of AMPAR-mediated synaptic currents obtained at each stimulation intensity decreased for DUP mice as compared to their littermates (Fig. 6A). Inhibitory interneurons are also present between thalamic terminals and LA neurons, forming a feedforward inhibition circuit. We evidenced a significant decrease of activation in the indirect recruitment of local interneurons in DUP mice compared to WT (Fig. 6B). Finally, we showed a significant decrease in AMPA/NMDA glutamate current ratio in DUP mice compared to WT (Fig. 6C). Taken together, these results indicate that synaptic function is impaired at thalamo-lateral amygdala synapses in DUP mice, in relation with a decrease of AMPA currents. These results are in full agreement with those reported in (35) and with an impaired AUTS2/AKT/mTORC1 pathway as reported in Fig. 2D.

**Fig. 6.**
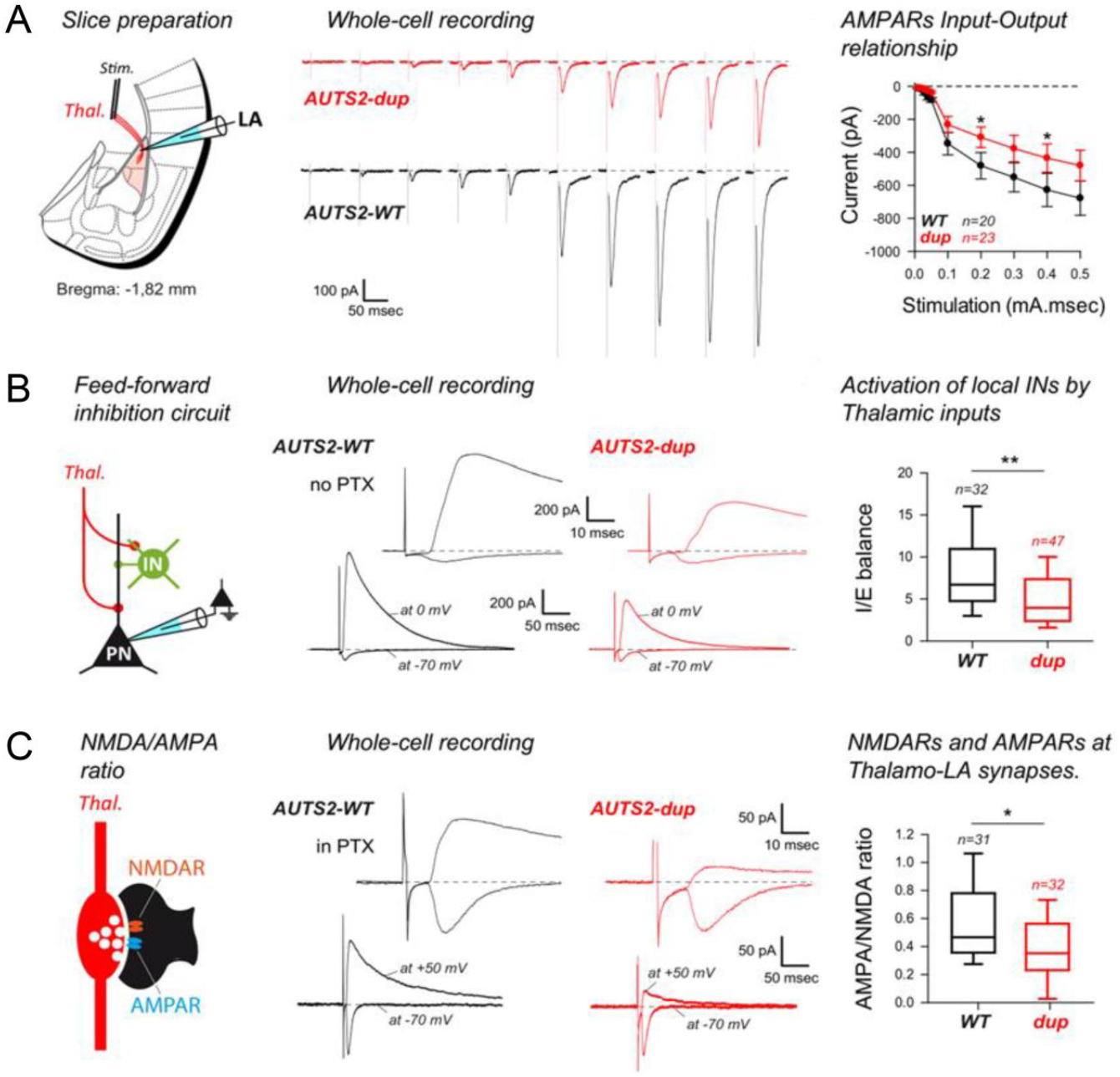
Analysis of thalamo-lateral amygdala synaptic function in Dup/+ Auts2 mice. **A**. Synaptic transmission at thalamo-lateral amygdala synapses was recorded in coronal acute slices of adult wild type (WT) and Auts2duplicated (DUP) mice (left). Excitatory transmission at thalamo-lateral amygdala synapses was evoked by electrical stimulations of increasing intensities delivered within the external capsule (middle). AMPAR-mediated synaptic currents obtained at each stimulation intensity were compared between WT and DUP mice. Number of recorded cells is indicated. *p<0.05. **B**. Feed-forward inhibition circuits in the lateral amygdala (left). AMPAR-EPSCs and GABAA-IPSCs were recorded in the same lateral amygdala principal cells respectively at −70mV and 0mV. The IPSC is observed after a significant delay (see insets) indicating the indirect recruitment of the local interneuron (middle). Comparison between the I/E balance observed following thalamic stimulations in lateral amygdala principal cells. Number of recorded cells is indicated. **p<0.01. **C.** Both NMDAR and AMPAR contribute to glutamatergic transmission at thalamo-lateral Amygdala synapses (left). In absence of GABAA inhibition (in PTX), NMDAR-mediated EPSCs can be recorded at +50mV. There, the residual current at 100msec after current onset is compared to the peak of current observed at −70mV (middle). AMPA/NMDA ratio were compared between WT and DUP preparations. Number of recorded cells is indicated. *p<0.05.

Molecular analysis of amygdala showed a significant decrease of the ratio phospho-Akt/Akt in DUP mice whereas no change was observed in temporal cortex, using western blotting (Fig. S9).

## Discussion

We were able to demonstrate that mouse Auts2 is not only localized in the nucleus but also in the dendritic spine, using synaptosomal fractions and super-resolution microscopy (Fig. 1-C). We also found that Aust2 directly interacts with the AKT E3 ligase TTC3 in the dendritic spine. Our study thereby provides a novel mechanistic and potentially therapeutic understanding of the mechanistic underpinnings of AUTS2 mutations, by demonstrating that AUTS2 regulates synaptic AKT/mTORC1 signaling and ubiquitination processes linked to AKT. Alterations in ubiquitin degradation has been previously associated with other de novo variants found in ASD patients (36). The importance of translational control in ASD is underscored by the number of single gene mutations in mTORC1 upstream regulators (PTEN, TSC1/TSC2, FMR1 or SHANK3) (37). In the first three cases, enhanced translation rates have been postulated to cause ASD (38). It is important to note that deletion of PTEN or TSC1/TSC2 not only activates mTORC1, but also alters mTORC2 activity (39).

We propose that decreased translation rates in *Auts2* down-regulated dendritic spines is responsible for the decrease in GriA1 and GriA2 AMPA receptor subunits and Grin2b NMDA receptor subunits. For SHANK3-deficient neurons, translation rates are considered as diminished with the SHANK3 deficiency. The proposed mechanism is a decrease in the degradation of CLK2 that increases protein phosphatase 2A (PP2A) activity, which reduces AKT activity. Consequently, protein synthesis decreases, leading to neuronal dysfunction (24, 40). In our study, we demonstrated that AUTS2 inhibits TTC3, an E3 ligase of AKT, whose function is to increase AKT polyubiquitination. We found that pAKT decrease both in down regulation of AUTS2 or when AUTS2 is duplicated. We also found that this effect is specific for mTORC1 pathway as p-GSKβ levels, a bona fide marker of mTORC2 pathway, were not modified.

Manipulation of this AKT/mTORC1 pathway is possible. CLK2 inhibition corrects impaired social motivation in Shank3 ^ΔC/ΔC^, by inhibition of PP2A that acts in the AKT/mTORC1 pathway (24, 40). Here, we found that restoring AKT2 levels in knock-down Auts2 neurons is sufficient to restore amplitude of synaptic responses.

The Auts2-deficient transgenic models that have been reported display abnormal social interaction and communication deficits (16, 41–43) but also recognition memory and cued-fear associative learning impairments (41, 42). Interestingly, for the Del/+ Auts2 mice, we did not find changes in social interaction and communication. We found stereotypies (rearing), reduced interest for objects (impairment in marble burying), resistance to change (Y maze) and abnormal recognition memory that are part of the set of core symptoms of ASDs. In contrast to mice with partial deletion of *Auts2* (41, 42), we did not find abnormal cued-fear associative memory in complete Del/+ model. In the Dup/+ model, we evidenced abnormal cued-fear associative memory, resistance to change (Y maze) and abnormal recognition memory. Both resistance to change (Y maze) and abnormal recognition memory appears to be common to our Del/+ and Dup/+ models. This result is in full agreement with the diagnosis of ASD found for patients displaying either a complete deletion or duplication of the 1.2 Mb AUTS2 locus (10).

A limited amount of information is available on neural circuits involved in behavioural traits found in mouse models of ASDs. Dysfunctions in fronto-striatal brain circuits have been characterized as responsible for excessive grooming found as stereotypies in Sapap3 mouse model (44). Engagement in social interaction was shown to depend on the activity in prelimbic cortex (PreL)-to-Tail-of-Striatum (TS) projection neurons in wild-type mice and was restored in Shank3 ^ΔC/ΔC^ mice by the chemogenetic activation of PreL/TS projection (45).

However, to the best of our knowledge, no ASD model was reported with both impairment of cued- and contextual-fear associative memory as we described in Fig. 5. Neural networks involved in these two types of associative memory have been characterized (33, 34, 46, 47). We were able to show that thalamo-lateral amygdala synapses were impaired in Dup/+ Auts2 mice with a decrease in AMPA-dependent synaptic currents (Fig. 6). Similar decrease was evidenced in down-regulated Auts2 neurons in culture, suggesting a similar mechanism at dendritic spines. Interestingly, it was demonstrated that both cued- and contextual-fear associative memory require surface diffusion of AMPA receptors (35, 48, 49). We also found a similar decrease in pAKT in both Dup/+ Auts2 and down-regulated Auts2 neurons. Further work is needed to dissect molecular mechanisms leading to a similar decrease in pAKT. Dup/+ Auts2 mice can be also a unique model to analyze molecular and cellular mechanism involved in the impairment of contextual fear associative memory.

Novel object recognition (NOR) involves medial temporal lobe processing (50). The ventral stream from the occipital lobe (visual stimuli) processes information about object recognition with relays in the perirhinal cortex and the lateral entorhinal cortex (51). The ventral occipito-temporal region is known to be involved in reading processes (52). NOR impairment can be considered as a proxy for abnormal reading processes as found in dyslexia. Interestingly, *AUTS2* locus was recently found associated to dyslexia using analysis of GWAS data for 51,800 adults self-reporting a dyslexia diagnosis and 1,087,070 controls (9).

In summary, our study highlighted the function of AUTS2 in the AKT/mTORC1 pathway in dendritic spines. Our findings might provide new insight into the developmental and neural circuit mechanisms involved in stereotypies, repetitive behaviors with restricted interests, resistance to change and abnormal recognition memory that are part form the second set of core symptoms of ASDs. Restoring normal levels of GriA1, Gria2 and Grin2a subunits at excitatory glutamatergic synapses may be instrumental for a personalized medicine approach in the treatment of ASDs.

## Acknowledgments

The authors wish to thank Dionne Swor and Megan Sribour for technical help. This work was partly supported by INSERM, the ANR06-neuro FRAXAmRNP, ANR NanoDiaMed, FP7-HEALTH AgedBrainSYSBIO (to MS), by grant MH77235 and MH080234 (to JAG), by CNRS, INSERM, University of Strasbourg (UDS), the “Centre Européen de Recherche en Biologie et en Médecine”, ANR-10-IDEX-0002-02, ANR-10-LABX-0030-INRT, ANR-10-INBS-07 PHENOMIN (to YHé) and ANR, Labex brain Bordeaux and Conseil regional Aquitaine (to YHu). AMLB was supported in part by Fondation pour la Recherche Médicale and Fondation Bettencourt-Schueller

The authors are grateful to the animal caretakers and services of the PHENOMIN-ICS, to the members of the research group, to the IGBMC laboratory for their helpful discussion and finally to animal caretakers of Pole in vivo de l’institut interdisciplinaire de neuroscience. This work has been supported by the National Centre for Scientific Research (The funders had no role in study design, data collection and analysis, decision to publish, or preparation of the manuscript).

## MATERIALS & METHODS

### Yeast Two-hybrid screen

Random primers were used to construct a cDNA library from human foetal brain mRNA (Invitrogen) into the pP6 plasmid derived from the original pACT2, using blunt end ligation of a SfiI linker. A 97 million independent clone library was obtained in *E. coli*. The DNA from this library was transformed into *S. cerevisiae* by classical lithium acetate protocol. Ten millions independent colonies were collected, pooled, and stored at −80°C as aliquot fractions of the same library. Two-hybrid screens using a cell-to-cell mating protocol was performed as previously described42. Full length Aust2 was cloned in lexA based pB27 bait vector and 50 million interactions have been tested and selected using His3 reporter. Performed by Hybrigenics S.A., Paris, France (http://www.hybrigenics.com/services.html).

### Primary cell, cell line cultures and transfection

We used mice of the C57BL6 strain. E17.5 mouse cortical and hippocampal neurons were at the same time dissociated enzymatically (0.25% trypsin, DNase), mechanically triturated with a flamed Pasteur pipette, and plated on 35mm dishes (8×10^5^ cortical cells and 4×10^5^ hippocampal cells per dish) coated with poly-DL-ornithine (Sigma), in DMEM (Invitrogen) supplemented with 10% foetal bovine serum. Four hours after plating, DMEM was replaced by Neurobasal^®^ medium (Invitrogen) supplemented with 2mM glutamine and 2% B27 (Invitrogen). Cells were transfected with constructs using Lipofectamine2000 (Invitrogen), as described by the manufacturer. SH-5YSY cell lines were cultured in DMEM supplemented with 10% of foetal bovine serum in 100mm dishes and were transfected close to confluence with Lipofectamine2000 (Invitrogen). For neuritic (axonal and dendritic) analysis, primary neuronal cultures were transfected after one or two days in culture and analyzed on day two or on days, three, four and five. For dendritic spines analysis, neurons were transfected after seven or eight days in culture and analyzed on day 21 or 22 respectively. All the analyses were carried out with the Wright Cell Imaging Facility plug-in in ImageJ as described in(*31*).

### Primary cell cultures and infection

We used mice of the OF1 strain. E15.5 mouse cortical and hippocampal neurons were treated as described earlier and plated on 100mm dishes or 24-wells plates (40×10^5^ cells per plate and 7×10^5^ cells per well). Hippocampal neurons were used for immunocytochemistry analysis and cortical cells were infected with AAV constructs according to 10^10^GU/well and analyzed 48-72hours later at DIV21 (24-wells plates were fixed and 100mm dishes were harvested for protein extraction).

### Plasmid constructs

To silence mouse *Auts2*, we used sh-RNA p-GIPZ vector (#RMM4431-98725112 for sh1 and #RMM4431-99010075 for sh2), as well as the control Non-Silencing sh-RNA (scrambled #RHS4346) from OpenBiosystems clone library (www.openbiosystems.com).

### AAV vectors production

rAAV production was carried out using the AAV Helper-Free system (Stratagene/Agilent Technologies). The target sequence was selected in the coding region of the mouse *Auts2* gene (NM_177047, position 3015): 5’-GAGACTCCTCCGTTAGTAA-3’. shRNA oligonucleotides were designed with the loop TTCAAGAGA and cloned under the control of mU6 promoter. The mU6-shRNA cassette was cloned into pAAV-MCS previously modified by adding a fluorescent eGFP under the control of the ubiquitous promoter CMV. A control pAAV-eGFP-shScramble expressing a shRNA-Scramble sequence 5’-GCGCTTAGCTGTAGGATTC-3’ that has no match *in silico* to the mouse genome was generated together with pAAV-eGFP-shAuts2. HEK293T/17 cells were triple cotransfected with a mixture of three plasmids: pAAV-eGFP-sh, pHelper and pAAV2/9 containing cap genes of AAV serotype 9 (kingly supplied by Dr. Julie Johnson, Penn Vector Core, USA). At 48h after transfection, cells were harvested, lysed by three freeze/thaw cycles in dry ice-ethanol and 37C baths, further treated with benzonase (100U/mL, Sigma-Aldrich) for 30mn at 37°C and clarified by centrifugation. Viral vectors were purified by lodixanol gradient ultracentrifugation, dialyzed in phosphate buffered saline, concentrated using centrifugal filter devices (Amicon) and stored at −80°C. The titer of AAV was estimated by quantitative real-time PCR and adjusted to 4.10^12^ GU/mL

### Stereotaxic injection, brain slices and dendritic spine analysis

Adult male mice were anesthetized by a cocktail of xylazine (10mg/kg) and ketamine (100mg/kg)) in physiological serum (NaCl 0.9%) with the analgesic buprenorphine (0,05mg/kg). and mounted on a stereotaxic stage (Stoelting). The cranium over the brain was exposed by a midline sagittal incision. A hole was made about 2.5 to 3mm caudal from lambda by a tip of forceps and a glass needle was placed in CA3 hippocampal region (Bregma 1.4;2;-2). Injection of 1μL of AAV was improved using autoclaved Hamilton gastight syringe and gauge steel needles, injector Ultra Micro Pump III with syringe pump controller (#SYS-MICRO4; World Precision Instruments). After the surgery, mice were kept on a heating pad until they recovered from the anesthesia and then they were returned to their home cages. Mice subjected to AAV injection were sacrificed 48hours after the AAV injection. Brains were dissected and frozen 1mn in isopentan over dry ice (−30°C) and conserved at −80°C. Frozen brains were coronally sectioned (40μm) using a cryostat (ThermoScientific Microm HM550). Sections were scanned using laser scanning confocal microscope (Leica SP5 from PICPEN imagery platform Centre de Psychiatrie et Neuroscience) with a resolution of 70nm/pixel. Image analysis was carried out with ImageJ software (Wayne Rasband, NIH) as described in(*31*).

### Immunocytochemistry and confocal microscopy

Cells were fixed by incubation in 4% paraformaldehyde in phosphate-buffered saline (PBS) for 20 min at room temperature, permeabilised by incubation for 10 min at room temperature in 0.3% Triton X-100 in PBS and blocked for 30 min at RT with 3% bovine serum albumin (BSA) and 0.1% Triton X-100 in PBS. Exception to this protocole was made for cell surface labelling for which only cell membrane proteins were labelled and no total proteins as above. In this particular case, the cells were blocked right after being fixed without any permeabilisation. After blocking, the cells were then incubated overnight at 4°C with primary antibodies (rabbit anti-AUTS2, 1:200, Atlas Antibodies; rabbit anti-GRIA1, 1:10, Calbiochem; mouse anti-GRIA2, 1:100, Millipore; mouse anti-GRIN2BB, 1:500, Neuromab; rabbit anti-HOMER, 1:200, Synaptic Systems; mouse anti-MUNC18, 1:200, BD Biosciences; rabbit anti-pAKT(Ser473), 1:50, rabbit anti-pGSK3β, 1:100, Cell Signaling; Cell Signaling; rabbit anti-panAKT, 1:400, Cell Signaling; mouse anti-PSD95, 1:250, Synaptic Systems; guinea-pig anti-SYNPO; 1:500, Synaptic Systems; goat anti-TTC3, 1:200, Santa Cruz clone E-20), diluted in the blocking solution. Cells were washed three times within PBS and incubated for 1h at room temperature with the secondary antibody (Alexa-488 anti-mouse and anti-rabbit IgG, 1:2,000, Invitrogen; CY3 anti-rabbit, anti-goat and anti-guinea-pig IgG, Jackson Laboratories, 1:2000, 1:1000 and 1:800 respectively; CY5 anti-mouse and anti-rabbit IgG, Jackson Laboratories, 1:1000) diluted in the blocking solution. Cells were incubated with Alexa Fluor-488 phalloidin (Life Technologies), washed three times within PBS, and mounted with Prolong Gold Mounting Medium (Invitrogen) or Mounting Medium with DAPI (DuoLink). Sections were scanned using laser scanning confocal microscope (Leica SP5 from PICPEN imagery platform Centre de Psychiatrie et Neuroscience) with a resolution of 70nm/pixel. Image analysis was carried out with ImageJ software (Wayne Rasband, NIH).

### High resolution

#### Sample Preparation for dSTORM nanoscopy

Cell cultures were fixed for 20 min in PBS1x, pH 7.4, containing 4% paraformaldehyde (PFA) and 1% sucrose, followed by three washing steps. Fiducial markers (TetraSpeck microspheres, 100 nm diameter, Invitrogen T7279) were attached to the coverslips after fixation (1:200 dilution) for 5min at room temperature (RT). For immuno-labelling of Auts2, fixed neurons were permeabilized with 0.3% Triton X-100 for 10 min at RT, then blocked with PSB1x containing 0.1% Triton X-100 and 3% BSA, and then labelled over night at 4°C with antibodies against rabbit anti-Auts2 (Atlas antibodies, 1:200) and guinea-pig anti-SYNPO (Synaptic Systems, 1:500), followed by Alexa Fluor 647 (Invitrogen, 1:250–500) secondary antibodies (1h at RT). Finally, Phalloïdine-Alexa488 (LifeTech, 1μl/cvs) was applied for 10min at RT for actin staining. dSTORM was conducted in PBS (pH 7.4), containing 10% glucose, 50 mM b-mercaptoethylamine, 0.5 mg/ml glucose oxidase, and 40 mg/ml catalase, degassed with N2(*32*).

#### dSTORM super-resolution imaging

Single-molecule imaging was carried out as described elsewhere(*33*) on an inverted Nikon Eclipse Ti microscope with a x100 oil-immersion objective (N.A. 1.49), an additional x1.5 lens, and an Andor iXon EMCCD camera (image pixel size, 107 nm), using specific lasers STORM of Alexa Fluor 647 (405 and 639 nm). Movies of 2.104 frames were acquired at frame rates of 50 ms with EMgain of 300. The z position was maintained during acquisition by a Nikon perfect focus system. Conventional fluorescence imaging was conducted with a mercury lamp and specific filter sets for the detection of Alexa 488 (excitation 485/20 nm, emission 525/30 nm).

#### Super-resolution image processing

Super-resolution image reconstruction was carried out using the software ThunderStorm(*34*, *35*). First, cross-correlation was used for drift correction. Then, data point with less than 500 photons and with a localisation precision higher than 20 nm were filtered out and finally, points with less than 5 neighbours within an area of 30 nm were also filtered out. Data rendering was performed using a 2D-gaussian of standard deviation equal to the localisation precision, with a final pixel size of 10 nm. Once Auts2 dSTORM super-resolution image was merged with actin conventional image, dendritic spines were manually selected. The number of Auts2 clusters per sub-region was counted and using the ImageJ plugin, we measured the spine area, and the spine length and head width (minimum and maximum Feret diameters).

### In situ proximity ligation assays (PLA) on dendritic spines

Cells at DIV21 were fixed by incubation in 4% paraformaldehyde in phosphate-buffered saline (PBS) for 20 min at room temperature and permeabilised by incubation for 10 min at room temperature in 0.3% Triton X-100 in PBS and PLA was performed according to the instructions of the manufacturer (DuoLink). Primary antibodies used were rabbit anti-AKT2, 1:50, Abcam; rabbit anti-AUTS2, 1:200, Atlas Antibodies; rabbit anti-panAKT, 1:400, Cell Signaling; goat anti-TOMM20, 1:100, Santa Cruz clone C20, goat anti-TTC3, 1:200, Santa Cruz clone E-20. Cells were incubated with Alexa Fluor-488 phalloidin (Life Technologies) before being washed within Buffer B (DuoLink) and mounted with Mounting Medium with DAPI (DuoLink). Sections were scanned using laser scanning confocal microscope (Leica SP5 from PICPEN imagery platform Centre de Psychiatrie et Neuroscience) with a resolution of 70nm/pixel. Image analysis was carried out with ImageJ software (Wayne Rasband, NIH).

### Statistical analysis

T-test were performed with Excel Software and KS-test (Kolmogorov-Smirnov) with web software (Kirkman, T.W. (1996) Statistics to Use. *http://www.physics.csbsju.edu/stats/* (2015).

#### Measurement of total dendrite glutamate receptor number versus surface receptor number

Immunocytochemistry with or without permeabilisation was performed as described above using anti-GRIN2B, anti-GRIA1 and anti-GRIA2 antibodies Immunocytochemistry with permeabilisation labelled total cell proteins; i.e. both membrane and intracellular proteins, whereas immunocytochemistry without permeabilisation labelled cell surface proteins by restrained labelling to membrane proteins only. Confocal microscopy scanning was then performed, as well as image and statistical analysis as described above.

### Mouse tissue protein extraction

Brain, cortex and hippocampus from either 3-4 months or embryonic (E18) OF1 mice were dissected and homogenized (pool of three animals) in MLB buffer (1%NP40, 100mM NaCl, 20mM Tris pH7.4 in PBS with 1x protease and phosphatase inhibitor cocktail) for 10 min on ice and centrifuged 15 min at 10,000g at 4°C. The supernatants were stored at −80°C until used and lysate protein concentration was determined using the DCTM Protein assay (Biorad).

### Preparation of synaptosomes and protein extraction

Cortex from 3-4 months mice brains were dissected and homogenized (pool of six animals) in H buffer (0.32M sucrose, 5mM Hepes 1M pH7.4, 1mM EDTA) using a glass potter. The homogenate was centrifuged at 800g for 7mn to remove nuclei and debris, the supernatant was centrifuged at 9,200g for 10mn to remove cytosolic supernatant. The pellet was resuspended in H buffer and gently stratified on a discontinuous Percoll gradient (5%, 10% and 23% v/v in H-buffered percoll) and centrifuged at 20,000g for 11mn. The layer between 10% and 23% Percoll (synaptosomal fraction) was collected and washed in H buffer by centrifugation. The synaptosomal pellets were resuspended in MLB buffer (1%NP40, 100mM NaCl, 20mM Tris pH7.4 in PBS with 1x protease and phosphatase inhibitor cocktail) for 10 min on ice and centrifuged 15 min at 10,000g at 4°C. The supernatants were stored at −80°C until used and lysate protein concentration was determined using the DCTM Protein assay (Biorad).

### Preparation of nuclear and cytosolic fraction and protein extraction

Cortex from 3-4 months mice brains were dissected and homogenized (pool of three animals) in nuclear and cytoplasmic reagents from the kit NE-PER as described by the manufacturer (Thermo Scientific).The cytoplasmic and nuclear supernatants were stored at −80°C until used and lysate protein concentration was determined using the DCTM Protein assay (Biorad).

### Cell line protein extraction

SH-5YSY cells transfected with shNS (n=5) and shD (n=5) were homogenised in RIPA buffer (0.5% NADOC, 1% NP40, 1% SDS 10% in PBS with 1x Protease Inhibitor Cocktail and 1% Phosphatase Inhibitor cocktails) for 10 min on ice and centrifuged 15 min at 10,000g at 4°C. The supernatants were stored at −80°C until used and lysate protein concentration was determined using the DCTM Protein assay (Biorad).

### Primary neuron protein extraction

Cortical primary neurons with AAV-Scramble (n=10) and AAV-shAuts2 (n=10) were homogenized for 10 min on ice in either MLB buffer (1%NP40, 100mM NaCl, 20mM Tris pH7.4 in PBS with 1x protease and phosphatase inhibitor cocktail) for AAV validation studies or in Triton buffer (0.15M NaCl, 50mM Tris pH7.4, 5mM EDTA, 1%Triton X-100 in PBS with 1x protease and phosphatase inhibitor cocktail) for molecular pathway studies and centrifuged 15 min at 10,000g at 4°C. The supernatants were stored at −80°C until used and lysate protein concentration was determined using the DCTM Protein assay (Biorad).

### Transgenic tissue protein extraction

Amygdala, hippocampus and cortex from 4 months transgenic mice were dissected and individually homogenized in Triton buffer (0.15M NaCl, 50mM Tris pH7.4, 5mM EDTA, 1%Triton X-100 in PBS with 1x protease and phosphatase inhibitor cocktail) for 10 min on ice and centrifuged 15 min at 10,000g at 4°C. The supernatants were stored at −80°C until used and lysate protein concentration was determined using the DCTM Protein assay (Biorad).

### Western blot analysis

Forty micrograms of protein was heated in the loading buffer and loaded on 5–12% gradient NuPage gels (Invitrogen). After the transfer on nitrocellulose membranes, membranes were incubated for 1h at room temperature in blocking solution (5% non-fat dried milk and 0.1% Tween 20 in PBS) and then overnight at 4°C with the primary antibody (mouse anti-β-ACTIN, 1:30,000, Sigma; rabbit anti-AKT1, 1:1000, Cell Signaling; rabbit anti-AKT2, 1:1000, Cell Signaling; rabbit anti-AKT3, 1:1000, Cell Signaling; rabbit anti-AUTS2, 1:1000, Abcam; rabbit anti-FBL, 1:1000, Cell Signaling; rabbit pAKT(Ser473), 1:500, Cell Signaling; rabbit anti-panAKT, 1:500, Cell Signaling; rabbit anti p-p70S6K(Ser371), 1:500, Cell Signaling; mouse anti-PSD95, 1:1000, Synaptic Systems). Membranes were washed in 0.1% Tween 20 PBS and incubated 1h at room temperature with the secondary antibody HRP-conjugated (anti-rabbit and anti-mouse IgG, 1:5,000, ECL) or the secondary fluorescent-conjugated (Dylight anti-rabbit 650 and anti-mouse 550 IgG, 1:5000, Fisher Scientific). Membranes were washed within 0.1% Tween 20 PBS. For HRP membranes, chemiluminescent detection reagents (BioRad) were used according to company instructions. Bands detection was performed with StainFree Chemidoc Imaging System (BioRad) and subjected to densitometric analysis using ImageLab software (BioRad).

### Primary cell sorting and total RNA extraction

E15.5 mouse telencephalic neurons were transfected at DIC1 and were washed with HBSS medium at DIC2, dissociated with a 0.05% trypsin, 0.05mM EDTA in HBSS and resuspended in DMEM supplemented with 10% fetal bovine serum (Invitrogen). GFP-positive cells were sorted using an Influx 500 (Cytopeia, Seattle, WA, USA) cell sorter as described: 2×14,000 GFP-positive cells from sh2-transfected cells (sh2) and 6,300 GFP-positive cells from sh1-tranfected cells (sh1). Using FACS, we also obtained in the same experiment the non-fluorescent fraction which was used as an internal control (NT): 3×14,000 GFP-negative cells from sh-transfected cells. For each group, cells were collected in XB buffer (Alphesys) for RNA extraction as described by the manufacturer. RNA quality was estimated using RNA pico 6000 Agilent chips.

### Primary cell infection and total RNA extraction

E15.5 mouse cortical neurons were infected at DIC4 and were washed with PBS medium at DIC6. Neurons were homogenized in Trizol reagent (Life Technologies) and RNA extraction was performed as described by the manufacturer and the treated with DNase I (Ambion). RNA quantity was estimated using Nandrop.

### Real-time quantitative RT-PCR analysis

Reverse transcription (RT) was carried out at 37°C for 50 min in a 20μl RT mixture containing 1°μg total RNA, 200U M-MLV reverse transcriptase (Invitrogen), 0.5mM of each dNTP and 6.25μM random hexamers (Invitrogen). The cDNAs generated were amplified by real-time PCR, using probes (obtained from PROLIGO Genset) labeled at their 5’ends with a fluorogenic reporter dye (FAM) and at their 3’ends with a quencher dye (TAMRA). Sequences of primers and probes are available on request. PCR assays had a final reaction volume of 20 μl, and contained 2U of Taq polymerase (Applied Biosystems), 10 μM primers and fluorogenic probe. PCR was carried out over 40 cycles of 95°C for 15 s, 60°C for 1 min and 50°C for 1 min. We used the Opticon2 sequence detection system (MJ Research/Biorad), with the Opticon Monitor software for data analysis. For each group, the cDNAs synthesized from total RNA were serially diluted to cover the 0.19 ng to 50 ng range for specific mRNAs and 0.98 pg to 250 pg range for 18S rRNAs. These serial dilutions were used to construct standard curves for 18S and for each gene of interest and to calculate the amounts of RNA for 18S and the genes of interest corresponding to the PCR products generated from inDICidual cDNAs for each experimental group. Each Q-PCR signal was normalized with respect to 18S and expressed in arbitrary units as in(*36*).

### Electrophysiological recordings of cortical neurons

7 to 10 days in vitro cortical neurons grown on coverslips in neurobasal medium supplemented with B27 were transfected either with sh-control (sh-NS) or sh-Auts2 (sh1) using lipofectamine (Invitrogen, France) according to manufacturer instructions. We used whole-cell patch-clamp recording of transfected neurons 48 to 72h after transfection. The cells were perfused voltage-clamped at –70 mV. The whole-cell with a Tyrode solution containing (in mM): 150 NaCl, 2 CaCl2, 1 MgCl2, 4 KCl, 10 glucose, 10 HEPES (pH 7.4). Spontaneous excitatory postsynaptic currents (sEPSCs) were studied in transfected cells voltage-clamped at –70mV. Patch electrodes, fabricated from thick borosilicate glass, were pulled and fire-polished to a final resistance of 3-4 MΩ and filled with the standard internal solution (in mM): 100 CsMES, 20 CsCl, 2 MgCl2, 5 ethylene glycol tetraacetic acid (EGTA), 10 HEPES, 4 ATP, and 15 phosphocreatine (pH 7.4).

#### Akt2 ubiquitination

HeLa cells were transfected with pCMV6-HA-Akt2, wild type human TTC3 in pCMV-Tag2 vector (Stratagene), mouse AUTS2-myc-DDK in pCMV6 (OriGene) and pCGN-His-Ub (His-tagged ubiquitin expression vector) by Lipofectamine LTX plus. Six teen hours after the transfection, the cells were treated with 10μM MG132 for additional 6hrs, rinsed twice with ice cold PBS, and lysed with Brij97 lysis buffer in the presence of proteinase inhibitors (leupeptin and AEBSF), 10μM MG132, and 5mM iodoacetamid. Immunoprecipitations were essentially performed as described in(*37*). The resulted cell lysates were precleaned with 50% slurry of ProA/ProG beads mixture for 30 minutes, immunoprecipitated with anti-HA antibody (12CA5, Roche) and the samples were resolved onto SDS-PAGE (6% Tris-Glycine Gel), immunoblotted by anti-HA (3F10, Roche) or Anti-His (Amersham), and detected by ECL. In the HA-blot, peroxidase-conjugated Anti-HA antibody (Boehringer-Mannheim) was utilized hence no immunoglobulin chains were visible. The expression of TTC3 and AUTS2 was detected by anti-Flag (M2, Sigma).

#### Brij97 cell lysis buffer

0.875% Brij97 (Sigma), 0.125% NP40, 150mM NaCl, 10mM Tris HCl pH 7.5, and 2.5 mM EDTA with proteinase inhibitor mix (Leupeptin and AEBSF), phosphatase inhibitors (1 mM Na3VO4 and 10 mM NaF) 10μM MG132, and 5mM iodoacetamide were added(*38*, *39*).

### Generation of the Aust2 deletion and duplication mutant lines

We decided to duplicate the whole genomic region containing the murine *Auts2* gene using the TAMERE strategy(*40*). The targeted region contains all the Auts2 isoforms also with the latest genomic annotations. A targeting construct was generated by PCR cloning of a 4.5 kb (5’ homology arm) and 3.5 kb (3’ homology arm) C57BL/6J genomic fragment in a vector containing a LoxP site and a Flipped NeoR selection marker. The linearized construct was electroporated in C57BL/6N mouse embryonic stem (ES) cells. After selection, targeted clones were identified by PCR using external primers and further confirmed by Southern blot with Neo and 3’ external probes. Two positive ES clones were injected into BALC/cN blastocysts, and male chimaeras derived gave germline transmission. The FRT surrounded selection marker was removed by breeding the chimeras with a Flp deleter line. One line was maintained and used for further breeding. This construct introduced a 5’ LoxP site 14914 bp at the 5’ side of Auts2. A second targeting construct was generated by PCR cloning of a 4 kb (5’ homology arm) and 3.1 kb (3’ homology arm) C57BL/6J genomic fragment in a vector containing a LoxP site and a F3 surrounded HygroR selection marker. The linearized construct was electroporated in C57BL/6N mouse embryonic stem (ES) cells. After selection, targeted clones were identified by PCR using external primers and further confirmed by Southern blot with Neo and 5’ external probes. Two positive ES clones were injected into BALC/cN blastocysts, and male chimaeras derived gave germline transmission. The F3 surrounded selection marker was removed by breeding the chimeras with a Flp deleter line. One line was maintained and used for further breeding. This construct introduced a 5’ LoxP site 12358 bp at the 3’ side of Auts2. The 3’ LoxP targeted line was bred with the *Hprt1^tm1(Cre)Mnn^* line(*41*), backcrossed on the C57BL/6N genetic background for more than 10 generation. The double heterozygotes mouses and the heterozygous 5’ LoxP mice were bred together in order to obtain females carrying LoxP sites on both homologous chromosomes and the Cre transgene. The carrier female mice was bred with wild-type C57BL/6N individual and the pups born from this breeding were PCR-genotyped for the deletion and duplication as this event is expected at the meiosis in the female germ line. The size of the deleted or duplicated fragment is 1.13 Mb. Hundred and fifty seven mice were genotyped by PCR. Five of them had the duplication of the whole *Auts2* locus and bared 2 the deletion of the whole gene. The sequence of both PCR products was confirmed by Sanger sequencing. We used the following for genotyping: Deletion (size of the PCR product 463 bp): EfA; GCAAACTCCAGAAAAGGCAGTACC with ErB; GCAGCGGCTGCAATTAACACAGA; Duplication (size of the PCR product 318 bp): EfB; TTCAGTGCCATGCTGGGAATAGAGC with ErA; CTTCCGGGTTAGGTTGTTTGTGG. The exact position of the deletion and the duplication on mouse chromosome 5 between 131,427,079 and 132,557,948 (GRCm38/mm10). The deletion Del(Auts2)1Ics and the corresponding duplication Dp(Auts2)2Ics, that will be noted here Del/+ and Dup/+, were established on a pure C57BL/6N genetic background.

### Behaviour analysis

Experimental procedures were approved by the local ethical committee Com’Eth under the accreditation number 2012-069. YH was the principal investigator of this study (accreditation 67-369). A series of behavioural experiments were conducted in order to evaluate cognition in the Dup/+ and Del/+ mice. For all these tests, mice were kept in SPF conditions with free access to food and water. The light cycle was controlled as 12 h light and 12 h dark (lights on at 7:00 AM) and the tests were conducted between 9:00 AM and 4:00 PM. To produce experimental groups, only animals coming from litters containing a minimum of two male pups were selected. After weaning, male mice were gathered by litters in the same cage. Animals were transferred to the experimental room 30 min before each experimental test. The tests were administered in the following order: Ymaze, open field, novel object recognition, sociability test, burying test, circadian activity and fear conditioning. Behavioural experimenters were blinded as to the genetic status of the animals.

#### Y maze

Short-term memory was assessed by recording spontaneous alternation in the Y-maze test(*42*). The Y-maze test is based on the innate preference of animals to explore an arm that has not been previously explored, a behaviour that, if occurred with a frequency greater than 50%, is called spontaneous alternation behavior (SAB). The maze was made of three enclosed plastic arms, set at an angle of 120° to each other, in the shape of a Y. The wall of the arm have different pattern to encourage SAB. Animals were placed at the end of one arm (this initial arm was alternated within the group of mice to prevent bias of arm placement), facing away from the center, and allowed to freely explore the apparatus for 8 min under moderate lighting conditions (70 lux in the center-most region). The time sequences of entries in the 3 arms were recorded. Alternation is determined from successive entries of the three arms on overlapping triplet sets in which three different arms are entered. The number of alternations is then divided by the number of alternation opportunities namely, total arm entries minus one. In addition, total entries were scored as an index of locomotor activity.

#### Open field (OF)

The open field test measures a combination of locomotor activity, exploratory drive and some aspects of anxiety and fear in mice(*43*). The output of the various interacting drives is locomotion, which is the direct measure obtained. The apparatus consisted of a white circular arena placed in a dimly lit testing room (20 lux). The mice were placed for 15 min in this arena. During this sessions, mice were monitored using a video tracking system (Ethovision, Wageningen, The Netherlands), total distance travelled in the different zone of the arena (center, intermediate and peripheral) was recorded.

#### Novel Object Recognition (NOR)

The novel object recognition task is based on the innate tendency of rodents to differentially explore novel objects over familiar ones(*44*). This test was done 24 hours after the OF session in the same apparatus. On Day 1, mice were free to explore 2 identical objects for 10 min. After this acquisition phase, mice returned to their home cage for a 24 hours retention interval. In order to test their memory, on day 2, one familiar object (i.e. already experienced during the acquisition phase) and one novel object were placed in the apparatus and mice were free to explore the two objects for a 10 min period. Between trials and subjects, the different objects were cleaned with 70° ethanol in order to reduce olfactory cues. To avoid a preference for one of the two objects, the new one is different for the different animal groups and counterbalanced between genotype. Is also the case for the emplacement of the novel object compared to the familiar one (left or right). Object exploration was manually scored and defined as the orientation of the nose to the object at a distance < 1 cm. For the retention phase, the percent of time exploring familiar vs novel objects was calculated to assess memory performance.

#### Marble burying test

This test is a species-typical behaviour that is responsive to many factors like hippocampal function, anxiety, depression or obsessive-compulsive disorder(*45*, *46*). Twenty glass marbles were placed, evenly spaced on a 5-cm layer of sawdust bedding which was lightly pressed down to make a flat even surface, in a plastic cage (of the type to which the mice had been housed). A mouse was placed in each cage and left for 20 min after which the number of marbles buried (to at least 2/3 their depth) with sawdust was counted.

#### Circadian activity

Circadian activity was measured to assess spontaneous activity and feeding behaviour over the complete light/dark cycle as described previously. Testing was performed in individual cages (11 x 21 x 18 cm^3^) fitted with infrared captors linked to an electronic interface (Imetronic, France) that provided automated measures of position and locomotor activity. Mice were put into cages at 6 pm on the first day and removed on the day 3 at 7 am. The light cycle was controlled as 12h light and 12h dark (lights turn on at 7 am). Feeding behaviour was evaluated using an automated lickometer and a 20 mg pellet feeder (Test Diet, Hoffman La-Roche).

#### Three-chambered social behavior test

The system, available from Stoelting (Dublin, Ireland), is composed of three successive identical chambers (20 x 40 x 22(height) cm) with 5 x 8cm openings allowing access between the chambers. The protocol was similar to the one described previously(*43*). During the habituation phase, the tested mouse was placed in the middle chamber and allowed to explore the three chambers under video tracking for 10 min, with each of the two side chambers containing an empty wire cage. The second phase of the test corresponding to the sociability test was carried out directly after the habituation phase. The test mouse was enclosed in the central box, while an unfamiliar mouse (stranger 1) was placed in one of the wire cages in a random manner. The doors were re-opened and the test mouse was allowed to explore the entire social test box for 10 min. Time spent sniffing each wire cage were recorded. The third phase tests is the preference for social novelty. A new stranger mouse (stranger 2) was placed into the empty wire cage and the test mouse was allowed to explore again the entire social test box for 6 min, having the choice between the first, already-known mouse (stranger 1) and the novel unfamiliar mouse (stranger 2). Stranger mice are adult C57BL/6J males (age 3-4 months) that were maintained in a different room from the tested mice to avoid olfactory and visual contacts. Several days before the test, stranger mice were habituated to the test in the wire cage 5 to 10 min per day for 5 days. Each stranger mouse was only used twice per test day and chosen randomly for either the sociability session or the preference for social novelty session.

#### Fear conditioning (Contextual and Cued)

To challenge hippocampus mediated cognitive behaviours we used the fear conditioning. This is an associative learning paradigm for measuring aversive learning and memory. In the fear conditioning procedure, a neutral conditioned stimulus (CS) such as light and tone is paired with an aversive unconditioned stimulus (US) such as mild foot shock. Concomitantly, animals associate the background context cues with the CS. After conditioning, the CS or the spatial context elicits a central state of fear in the absence of the US, expressed as reduced locomotors activity or total lack of movement (freezing). The Immobility time is used as a measure of learning/memory performances(*47*, *48*). Experiments were conducted in four operant chambers (28 × 21 × 22 cm) with a metal bar floor linked to a shocker (Coulbourn Instruments, Allentown US). Chambers were dimly lit with a permanent house-light and equipped with a speaker for tone delivery and an infra-red activity monitor. The activity/inactivity behaviour was monitored continuously during the different session and data were expressed in duration of inactivity per 2 s. The experimental procedure comprises 3 sessions over 2 days. In day 1, for the conditioning session, the mouse was allowed to acclimate for 4 min, then a light/tone (10 kHz, 80-dB) CS was presented for 20s and terminated by a mild foot shock US (1sec, 0.4 mA). After the foot shock, animals were left in the chamber for another 2 minutes. We defined total freezing time in first 2min and 4min and 2min immediately after foot shock as PRE1 PRE2 and POST, respectively. In Day 2, the context testing was performed by placed back the mouse into the same chamber and allowed to explore for 6 minutes without presentation of the light/auditory CS. The movement of the animal was monitored to detect freezing behaviour consequent to recognition of the chamber as the spatial context (contextual learning). The total freezing time is calculated by 2min block as CONT2, CONT4 and CONT6. Finally, the cue testing was performed 5 hours after the context testing. Animals were tested in modified conditioning chambers with walls and floor of a different colour and texture. The mouse was allowed to habituate for 2 minutes then presented to light/auditory cues for 2 minutes to evaluated conditioning fear. The total freezing time was calculated by 2min block as PRECUE1, CUE1, PRECUE2 and CUE2.

### *Electrophysiological recordings of transgenic mice* (as in (*49*))

Ethics Statement: Every effort was made to minimize the number of animals used and their suffering. The experimental design and all procedures were in accordance with the European guide for the care and use of laboratory animals and the animal care guidelines issued by the animal experimental committee of Bordeaux Universities (CE50) (A5012009). All experiments were performed using male Auts2 x/x (Auts2 KI) and their control x/x (Auts2 WT) littermates (C57BL/6J background, 2-4 months old), housed in 12/12 LD with ad libitum feeding.

#### Acute slices preparations

Due to animal age (3-4 months old), we used the “protective recovery method” described in http://www.brainslicemethods.com. Briefly, mice were anesthetized with a mixture of ketamine/xylazine (100mg/kg and 10mg/kg respectively) and cardiac-perfused with ice-cold, oxygenated (95% O2, 5% CO2) cutting solution (NMDG) containing (in mM): 93 NMDG, 93 HCl, 2.5 KCl, 1.2 NaH2PO4, 30 NaHCO3, 25 Glucose, 10 MgSO4, 0.5 CaCl2, 5 Sodium Ascorbate, 3 Sodium Pyruvate, 2 Thiourea and 12mM N-Acetyl-L-cystéïne (pH 7.3-7.4, with osmolarity of 300-310 mOsm). The brains were rapidly removed and placed in the ice-cold and oxygenated NMDG cutting solution described above. Coronal slices (300 μm) were prepared using a Vibratome (VT1200S, Leica Microsystems, USA) and transferred to an incubation chamber held at 32°C and containing the same NMDG cutting solution. After this incubation the slices were maintained at room temperature in oxygenated modified ACSF containing (mM): 92 NaCl, 2.5 KCl, 1.2 NaH2PO4, 30 NaHCO3, 20 HEPES, 25 Glucose, 2 MgSO4, 2 CaCl2, 5 Sodium Ascorbate, 3 Sodium Pyruvate, 2 Thiourea and 12mM N-Acetyl-L-cystéïne (pH 7.3-7.4, with osmolarity of 300-310 mOsm) until recording.

#### Recordings

Whole-cell recordings from BLA principal neurons were performed at 30-32°C in a superfusing chamber as previously described (Houbaert et al. 2013, JNS). Neurons were visually identified with infrared videomicroscopy using an upright microscope equipped with a 60x objective. Patch electrodes (3-5 MΩ) were pulled from borosilicate glass tubing and filled with a low-chloride solution containing (in mM): 140 Cs-methylsulfonate, 5 QX314-Cl, 10 HEPES, 10 phosphocreatine, 4 Mg-ATP, and 0.3 Na-GTP (pH adjusted to 7.25 with CsOH, 295 mOsm). LA principal neurons were recorded in voltage clamp mode at −70 mV (to record AMPAR-mediated EPSCs), 0 mV (to record GABAAR-mediated IPSCs), or +50 mV (NMDAR mediated EPSCs) as previously described(*50*). Cortico-LA monosynaptic EPSCs and di-synaptic IPSCs were elicited by 1 msec electrical stimulations delivered by a bipolar electrode positioned in the external capsule, connected to an external, isolated stimulator.

#### Data acquisition and analysis

Data were recorded with a Multiclamp700B (Molecular Devices, USA), filtered at 2 kHz and digitized at 10 kHz. Data were acquired and analysed with pClamp10.2 (Molecular Devices).

